# Measuring X inactivation skew for retinal diseases with adaptive nanopore sequencing

**DOI:** 10.1101/2024.03.20.585856

**Authors:** Sena A Gocuk, James Lancaster, Shian Su, Jasleen K Jolly, Thomas L Edwards, Doron G Hickey, Matthew E Ritchie, Marnie E Blewitt, Lauren N Ayton, Quentin Gouil

**Affiliations:** Department of Optometry and Vision Sciences, The University of Melbourne, Parkville Victoria, Australia; Centre for Eye Research Australia, East Melbourne, Victoria, Australia; Ophthalmology, Department of Surgery, University of Melbourne, Melbourne, Victoria, Australia; Epigenetics and Development Division, The Walter and Eliza Hall Institute of Medical Research, Parkville, Victoria, Australia; Department of Medical Biology, The University of Melbourne, Melbourne, Victoria, Australia; Vision and Eye Research Institute, Anglia Ruskin University, Cambridge, United Kingdom

**Author notes:** These authors contributed equally as co-first (Sena A Gocuk, James Lancaster). These authors contributed equally as co-last (Lauren N Ayton, Quentin Gouil).

**Keywords:** X-linked diseases, retinal disorders, skewed X inactivation, long-read sequencing, DNA methylation

## Abstract

X-linked genetic disorders typically affect females less severely than males due to the presence of a second X chromosome not carrying the deleterious variant. However, the phenotypic expression in females is highly variable, which may be explained by an allelic skew in X chromosome inactivation. Accurate measurement of X inactivation skew is crucial to understand and predict disease phenotype in carrier females, with prediction especially relevant for degenerative conditions.

We propose a novel approach using nanopore sequencing to quantify skewed X inactivation accurately. By phasing sequence variants and methylation patterns, this single assay reveals the disease variant, X inactivation skew, its directionality, and is applicable to all patients and X-linked variants. Enrichment of X-chromosome reads through adaptive sampling enhances cost-efficiency. Our study includes a cohort of 16 X-linked variant carrier females affected by two X-linked inherited retinal diseases: choroideremia and *RPGR*-associated retinitis pigmentosa. As retinal DNA cannot be readily obtained, we instead determine the skew from peripheral samples (blood, saliva and buccal mucosa), and correlate it to phenotypic outcomes. This revealed a strong correlation between X inactivation skew and disease presentation, confirming the value in performing this assay and its potential as a way to prioritise patients for early intervention, such as gene therapy currently in clinical trials for these conditions.

Our method of assessing skewed X inactivation is applicable to all long-read genomic datasets, providing insights into disease risk and severity and aiding in the development of individualised strategies for X-linked variant carrier females.

## Introduction

X chromosome inactivation (XCI) is a dosage compensation mechanism equalising the expression of the majority of X-linked genes between males and females [1]. In 46,XX females, one X chromosome is chosen stochastically as the inactive X (Xi) in a small number of cells early in embryonic development, and remains silenced throughout life [2]. Females are thus mosaics of cells where either one of the two X chromosomes is inactive. Skewed X inactivation, whereby allelic Xi status in a pool of cells departs from the predicted 50:50 ratio, may explain the phenotypic variability in female carriers of X-linked conditions, when the gene in question is subject to X inactivation [3]. Skew may be organism-wide (global skew observable in all or most tissues if the skew is due to biased X choice early in development), tissue-specific (when a skewed pool of cells give rise to a tissue, a common occurrence in highly clonal tissues like blood), or geographic (spatial heterogeneity within a tissue due to ontological patches of cells with the same Xi, like the coat of calico cats). XCI skew may be protective when the disease-causing variant is on the preferentially silenced X, or deleterious when on the preferentially active X. Skewed X inactivation is common, with estimates that 50% of females display skews of 65:35 or greater in blood [4, 5].

X-linked inherited retinal diseases, such as choroideremia and X-linked retinitis pigmentosa, most strongly affect males, with night blindness and peripheral vision loss during early adolescence which progress to central vision loss at later stages of the condition. Female carriers of X-linked inherited retinal disorders present with a spectrum of disease severities, ranging from near normal retinae to severe retinal degeneration, the latter known as “male-pattern” retinal degeneration [6, 7]. This variation and the observed prevalences is compatible with skewed X inactivation as a major factor of disease severity in females [8]. Retinal phenotypes of female carriers can present as a mosaic pattern comprising regional patches of degeneration due to retinal mottling and/or pigmentary changes, which would again be compatible with a geographic XCI skew. Although it would be ideal to investigate the role of X inactivation in inherited retinal disorders directly in the affected retinal epithelium and photoreceptor cells, retinal biopsy from living patients is not justifiable due to the risk of vision loss directly and indirectly relating to the procedure, as well as the costs and resources involved in obtaining samples [7].

To date, the evidence for the importance of skewed XCI for X-linked inherited retinal disorders remains limited [9]. As a proxy for the retina, X inactivation skew of female carriers of X-linked inherited retinal disorders has been previously reported in blood samples, based on DNA methylation measurements. This works because the inactive X chromo-some has hypermethylated CpG islands [10]. These studies used conventional methods based on methylation-sensitive restriction enzymes and PCR fragment length analysis [9, 11]. However, such methods provide incomplete information due to the lack of informative polymorphisms found in over 20% of individuals [12], quantitatively-limited analysis of just a few CpG sites and one or two genomic loci with the potential for PCR bias, the requirement for parental DNA to determine the origin of the variant and direction of the skew and additional genetic testing to ascertain the deleterious variant. A sequencing-based version of this test has recently been proposed [13], directly reading the methylation of the *AR* and *RP2* CpG islands by nanopore sequencing of Cas9-targeted native DNA. DNA modifications trigger recognisable variation in the electrical current measured at the nanopore, enabling base-level modification analysis in long reads. Though this targeted test improves quantitation accuracy, it suffers from most of the same limitations outlined above. Furthermore, blood samples have been found to increase in XCI skew with age and may therefore not accurately reflect the XCI status of the retina and other tissues [14, 15].

We set out to develop a new method to measure X inactivation skew based on long-read whole-genome sequencing, to simultaneously identify variants and investigate the contribution of skew to X-linked diseases in females. We used the Oxford Nanopore Technologies PromethION 24 platform and its adaptive sampling capacity to enrich X chromosome reads. We sequenced peripheral samples (blood, saliva and buccal mucosa) for a cohort of female carriers of X-linked inherited retinal disorders (choroideremia and *RPGR*-associated X-linked retinitis pigmentosa) to determine X inactivation skew in these tissues, infer the retinal skew and correlate with disease severity and retinal phenotype. We also include a female individual who did not have an inherited retinal disorder but for whom we were able to retrieve fresh retinal tissue (following enucleation for an invasive orbital tumour), allowing a direct comparison of skew in the retina and peripheral samples. Recent advances in the development of gene therapy for retinal diseases in males have sparked interest to determine whether female carriers, particularly those at risk of severe disease, should be considered for upcoming therapeutic intervention. Accurate measurement of skewed X inactivation may assist in determining the most suitable candidates.

## Results

### Optimising X chromosome enrichment with adaptive sampling

In order to increase the number of samples that could be run on one PromethION flow cell while still reaching a target coverage of 15–20X per sample, we made use of the adaptive sampling capacity of the PromethION P24 (beta feature in MinKNOW, where the sequencing of off-target reads is interrupted by voltage reversal to make the pore available for another fragment, Fig. 1A). To guide protocol optimisation, we estimated the relationship between read length and target coverage (see Methods). When targeting the entire X chromosome (154 Mb), maximising read length is beneficial, up to the point where pore blocking and decreased pore occupancy (longer fragments mean fewer molecules) become the dominant factors in reducing the output.

**Fig. 1.**
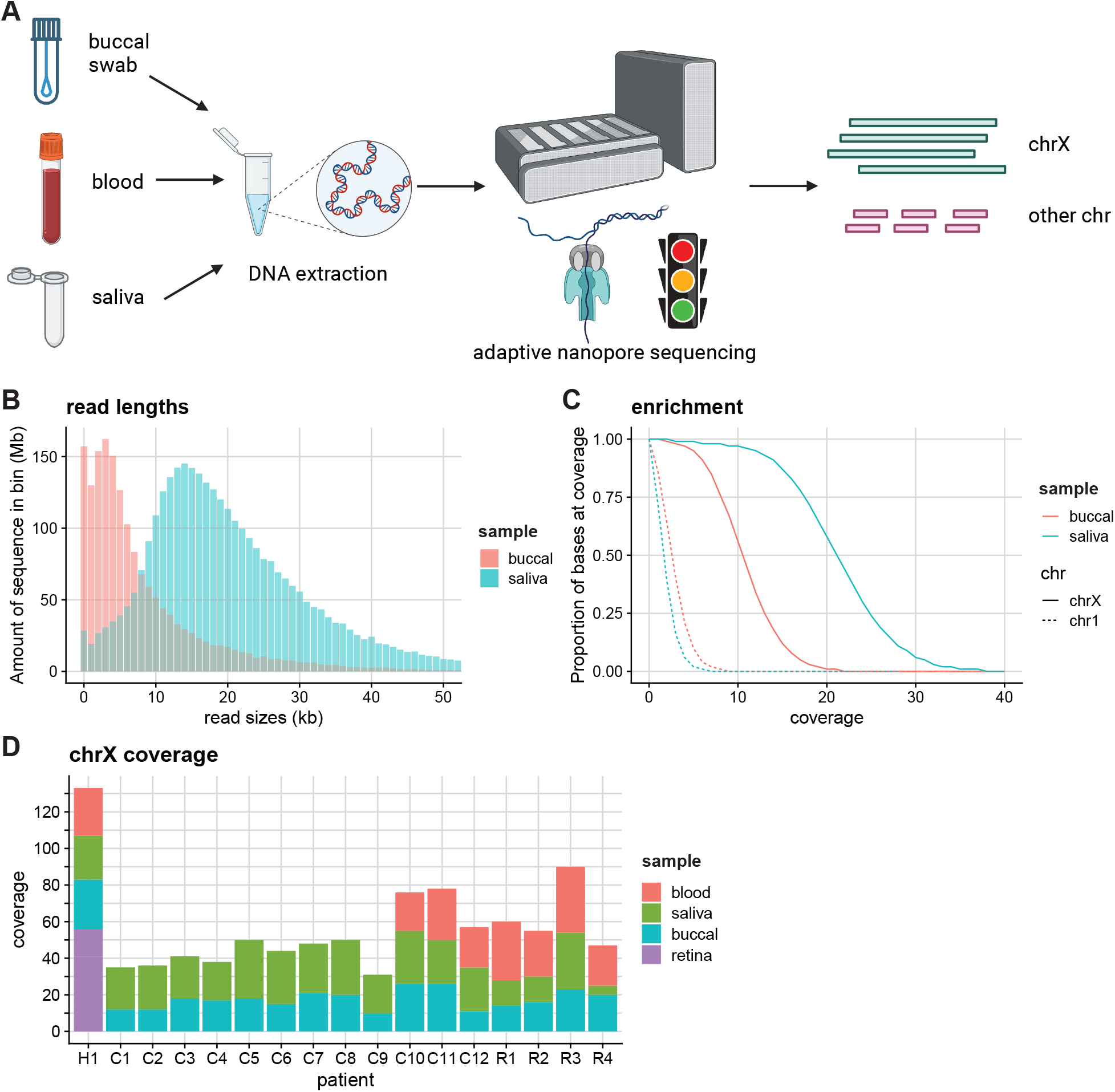
X inactivation skew sequencing workflow. **A)** Adaptive nanopore sequencing of peripheral samples (blood, saliva, and buccal mucosa) to maximise coverage of the X chromosome. Sequencing of fragments that do not map to the X chromosome is terminated early. **B)** Degraded buccal DNA results in shorter read lengths and **C)** reduced X chromosome enrichment performance. The read lengths for (accepted) reads mapping to the X chromosome are plotted for patient C9 for illustrative purposes. Rejected reads are typically under 1 kb. **D)** X chromosome coverage for each patient, split by sample. At least 30X combined X chromosome coverage was obtained for each patient. H1: healthy individual; C1-C12: female carriers of choroideremia; R1-R4: female carriers of *RPGR*-associated X-linked retinitis pigmentosa.

Our sequencing results corroborated this estimate, as libraries dominated by small fragments enriched poorly for X chromosome reads (Fig. 1B, C). Two factors contributed to small fragments dominating the sequencing: either the DNA samples were degraded (buccal swabs had consistently worse DNA integrity than blood or saliva) or in intact samples the unsheared, high-molecular weight fragments could not be efficiently sequenced. Therefore, we strived to increase the average fragment lengths in the libraries by a combination of clean-ups (selective precipitation when DNA amounts allowed, bead-based clean-ups in the case of low input DNA) and shearing to 10–20 kb fragments (Fig. 1B). With optimised libraries, we obtained up to 90X coverage of the X chromosome per PromethION flow cell, equivalent to a 3–4-fold enrichment over non-adaptive sequencing. We obtained at least 30X total coverage of the X chromosome for each patient, across 2–4 samples per patient, and providing sufficient data for analysis of alleles and DNA methylation.

### Calculating X inactivation skew by phasing alleles and epialleles

Next we developed a method using the phasing of long reads into alleles (maternal and paternal, or haplotype 1 and haplotype 2 when the parent-of-origin is unknown) and their clustering based on DNA methylation (Xi being hypermethylated and Xa being hypomethylated) to determine skewed X inactivation (Figure 2A). We generated a dataset where the maternal and paternal alleles are distinct by crossing two different mouse strains, and where X inactivation is genetically skewed through the inheritance of a mutated *Xist* [16, 17]. We made neural stem cells from the F1 animals and sequenced the DNA by nanopore according to the ultra-long sequencing protocol [17].

**Fig. 2.**
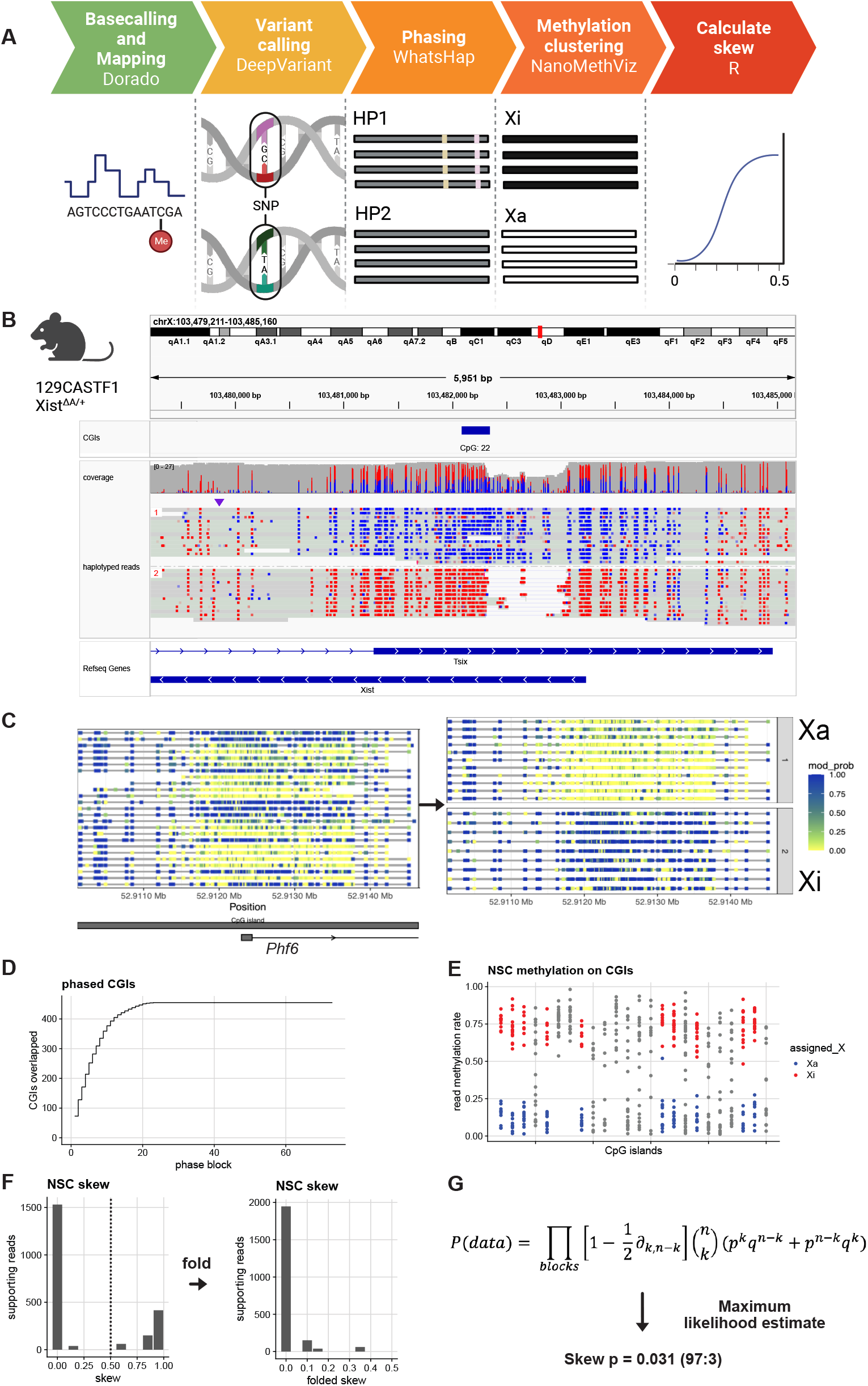
Bioinformatic workflow to measure X inactivation skew with long-read genomics, demonstrated on ground-truth data. **A)** Five-step bioinformatic pipeline: basecalling and mapping reads including modified bases, SNV calling, phasing alleles and reads based on SNVs, clustering reads based on their methylation profile, tabulating the allele/epiallele combinations to calculate the skew. **B)** Validation on mouse data with genetically engineered skewed X inactivation: a deletion of the *Xist* A-repeat (haplotype 2) triggers its constitutive choice as the Xi, as confirmed by the DNA methylation profile of the *Xist* promoter (red: methylated CpG; blue: unmethylated CpG). **C)** Example of read clustering by methylation profile for the *Phf6* CpG island: reads split into two clusters, one with low methylation (yellow, assigned as Xa) and one with high methylation (blue, assigned as Xi). **D)** Cumulative sum of CpG islands overlapped by haplotype blocks. Large haplotype blocks overlap many CpG islands, providing additional power to estimate the X inactivation skew in those haplotype blocks. **E)** CpG island methylation clustering results for one haplotype block. Each point represents a read. CpG islands with grey reads failed to split into Xi/Xa groups. **F)** Non-folded and folded histograms of the haplotype block-wise skews, binned in 10% bins and scaled by the underlying number of reads. Skew is measured as the proportion of reads supporting haplotype 1 as Xa. Because skewed X inactivation is genetically engineered in this experiment, haplotype blocks are expected to yield skews close to either 0 or 1. In the case of perfectly balanced X inactivation, haplotype blocks should return skews of 0.5. **G)** From the block skews, the sample skew *p* is calculated by maximising the joint probability of observing the data. *q* = 1 *− p*; for each haplotype block, *k* is the number of haplotype 1 reads that are Xa and haplotype 2 reads that are Xi (‘successes’) and *n* the total number of haplotyped and clustered reads (‘trials’).

After small variant calling with DeepVariant [18] and read haplotyping with WhatsHap [19], visualisation of the *Xist* locus confirmed the correct SNV-based separation of haplotypes with the presence of the *Xist*^ΔA^ mutation as a *∼*750-base deletion *∼*120 nucleotides downstream of the *Xist* transcriptional start site on haplotype 2 (from the 129/Sv strain background). It also illustrated the contrasting DNA methylation patterns between the active and inactive X, correlating with the *Xist* gene expression: highly methylated haplotype 2 with mCpGs in red (silenced, non-functional *Xist*), and methylation-depleted haplotype 1 with CpGs in blue, expressing a functional *Xist* (Fig. 2B). Of note, the methylation pattern at the *Xist* CpG island is reversed compared to the rest of the X chromosome CpG islands: the inactive X, which expresses *Xist*, is unmethylated (haplotype 1) while the active X (*Xist* silenced) is the methylated allele (haplotype 2). For all other CpG islands on the X chromosome, methylation-based clustering of reads assigns Xa to the group of low-methylation reads and Xi to the group of high-methylation reads (Fig. 2C). This clustering is performed using the density-based clustering algorithm DBSCAN within NanoMethViz [20]. The promoter CpG island of the X-linked *Phf6* gene illustrated how reads clustered in two well-separated groups (Fig. 2C).

Independent of their methylation profile, reads were also assigned to either haplotype 1 or haplotype 2. Regions of low SNV density or regions that are highly repetitive break up the haplotype phasing into haplotype blocks. As a result, haplotype 1 in one block may be the paternal allele while haplotype 1 in the next block may be the maternal allele (phase switching). Haplotype blocks are separated by genomic segments that cannot be haplotyped. In the mouse neural stem cell dataset, out of 469 annotated CpG islands (UCSC mm10 annotation), where we can expect DNA methylation on the inactive X, 454 were haplotyped (Fig. 2D), allowing the assignment of reads to haplotype 1 and haplotype 2.

Of these haplotyped CpG islands, 105 CpG islands clustered into two methylation-level clusters, amounting to 2,300 reads assigned as Xa or Xi (Fig. 2E). The remaining haplotyped CpG islands were not split into Xi and Xa methylation patterns for one of several reasons: similarly high or low methylation across all reads, lack of separation due to continuous variation in methylation, or insufficient read numbers to form a cluster (minimum set as 5 reads, Fig. 2E). We then tallied up the reads that were both haplotypes and clustered as Xa/Xi to calculate the X inactivation skew for each haplotype block. Large haplotype blocks may overlap multiple CpG islands, which allows pooling the counts of haplotype 1/haplotype 2 and Xa/Xi combinations for all CpG islands across the block. The haplotype block overlapping the most CpG islands covered 80 CpG islands (Fig. 2D). The haplotype-block skew for each block *i* is expressed as:

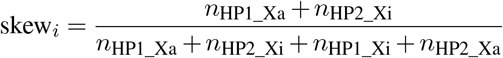

This represents the bias for haplotype 1 to be the active allele. Since phase switching may occur between blocks, we fold the skew: folded_skew_*i*_ = *min*(skew_*i*_, 1 *−* skew_*i*_), equivalent to mirroring the skew that is above 0.5 down to below 0.5 with the symmetry axis x = 0.5 (Fig. 2F). As expected in the genetically skewed mouse NSC dataset, skews for the haplotype blocks distributed around the extremes at 0 and 1, and the folded distribution peaked around 0 (Fig. 2F).

The global skew *p* for the sample is derived from the maximum likelihood estimate of the observed read counts in each block under the folded binomial model (Fig. 2G, [21, 22]). The numerical optimisation for *p* for the mouse NSC data yielded a global skew of 0.031, or a 97:3 ratio al-lele1_Xa:allele2_Xi, close to the expected 100:0.

### Measuring X inactivation skew in human retinal and peripheral samples

Having established the skew analysis pipeline on ground truth data, we next applied it to a patient for whom we could collect retina, blood, saliva and buccal swab samples. This allowed us to investigate the correlation in skew between tissues, as well as the spatial heterogeneity of the retina by collecting discs across four quadrants of the retina (Fig. 3A).

**Fig. 3.**
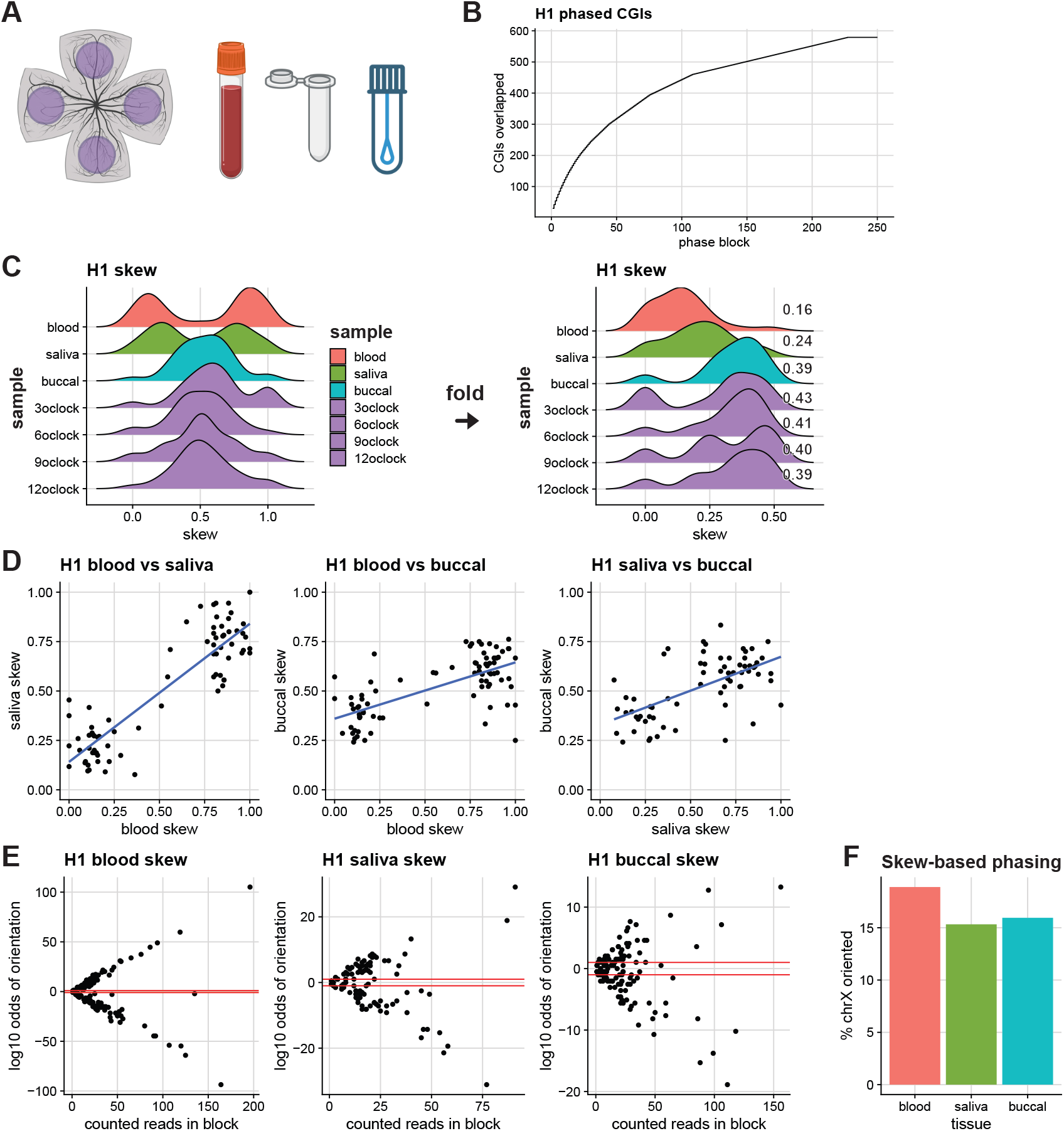
X inactivation skew across blood, saliva, buccal swab and retina in a healthy patient. **A)** Collected samples: 4 quadrants of the retina from one eye, blood, saliva and buccal swab. **B)** Cumulative sum of CpG islands overlapped by haplotype blocks. **C)** Non-folded and folded distributions of haplotype blocks’ skews in the seven samples. Blood and saliva show a clear skew, whereas buccal swab and all four retina samples are not highly skewed. **D)** Positive correlation of non-folded haplotype blocks’ skews across samples shows that the Xa allelic bias is consistent across tissues (the same X chromosome in the preferentially active X in all skewed tissues). **E)** Skewed X inactivation allows scaffolding haplotype blocks into alleles. Log-likelihood ratio of haplotype 1 being the preferential Xa over Xi for each block. A red line is plotted at y=1 (haplotype 1 ten times more likely than haplotype 2 to be the preferential Xa) and y=-1 (haplotype 2 ten times more likely than haplotype 1 to be the preferential Xa). **F)** Up to 18% of the X chromosome for patient H1 can be arranged in a consistent haplotype (higher-order phasing) based on haplotype block skew.

The 10–20 kb reads optimised for adaptive sampling of the X chromosome resulted in shorter haplotype blocks than with the ultra-long read mouse data, but overall 579 CpG islands (of 879 CpG islands) were phased into alleles (Fig. 3B).

The distribution of block skews revealed a strong X inactivation skew in blood (*p*_blood_ = 0.16), a slightly smaller skew in saliva (*p*_saliva_ = 0.24) and a more balanced X inactivation profile in the buccal swab and retinal discs (0.39 to 0.43, Fig. 3C). Each quadrant of the retina had a similarly balanced X inactivation profile, suggesting a well-mixed pattern of cells with the maternal or paternal X inactivated. The lack of correlation between the blood and retinal skews cautions against the ready extrapolation of retinal skew from blood skew, although it is common practice. Here the sampling of multiple tissues enabled us to rule out an organism-wide (global) skew in favour of a tissue-specific skew.

Among the samples collected, the buccal swab is the most likely to correlate with the retina. It is less susceptible to high clonal expansion than the cells present in blood and saliva [4, 23]. Although it is more challenging to extract high-quality DNA from it, the buccal swab may be the most informative for evaluating skew in the retina. In this particular patient’s case, it was the one that most closely matched the non-skewed retinal X inactivation profile.

### Using phase to orient skew and using skew to phase

Although from block to block the identity of haplotypes can switch, for a single patient the haplotype for a block is the same across tissues. Therefore the correlation in unfolded skew between blocks informs on the direction of the skew: a positive correlation indicates the skew is in the same direction in both samples, whereas a negative correlation would indicate a reversal of skew between samples.

We observed a strong positive correlation between the blood and saliva block skews (Fig. 3D), indicating that the skew direction was the same in both highly skewed samples. The correlation with the more mildly skewed buccal swab sample remained positive, again pointing to the same direction of skew.

We then asked whether we could improve phasing of the X chromosome by using the skew information: for a highly skewed sample, if in one block haplotype 1 is the predominantly active allele and in the next block haplotype 2 is the predominantly active allele, this would indicate that there was a phase switch between these two blocks. Even though the haplotype blocks remain disjointed, they could be oriented so that haplotype 1 is consistently the same parental haplotype across haplotype blocks (higher-order phasing). Strong skews gave very high certainty in orienting blocks (Fig. 3E), but even for the buccal sample around 16% of the X chromosome could be phased across haplotype blocks by using skew. Orienting the skew in each block is also helpful to assess whether a particular variant is likely to be on the predominantly active or inactive X.

Intriguingly, one haplotype block with a high number of phased and clustered reads could not be confidently oriented (outlier in Fig. 3E, blood panel). Further inspection revealed that this was due to an error in variant phasing in the difficult region harbouring duplicated *RHOXF2* and *RHOXF2B*: ambiguous read mapping over a 48.9-kb segmental duplication [24] led to a phase switch within the haplotype block (Supp Fig. 1). Haplotype 1 was predominantly the Xa upstream of the ambiguous region, whereas it became predominantly the Xi downstream. Thus the skew analysis enabled the identification and correction of a variant phasing error across this complex region.

### X inactivation skew in female carriers of inherited retinal disorders

Next we applied our method of measuring X inactivation skew to a cohort of patients with inherited retinal disorders: 12 carrying variants in the *CHM* gene causing choroideremia and 4 carrying variants in the *RPGR* gene, a major cause of X-linked retinitis pigmentosa (XLRP, Table 1). There were two patient pairs carrying the same variants: a mother and daughter pair, and two sisters. Seven carriers had grade 4 male-pattern degeneration (four with variants in the *RPGR* gene and three with variants in the *CHM* gene). All other carriers had variants in the *CHM* gene and had relatively milder retinal phenotypes: one carrier with grade 1 (fine, near normal), four with grade 2 (coarse), and four with grade 3 (geographic) [6].

**Table 1.** Variants in affected patients. Coordinates are relative to hg38 chrX (NC_000023.11), *CHM* MANE transcript NM_000390.4 and translated protein NP_000381.1, *RPGR* transcript NM_001034853.1 and protein NP_001030025.1. Patient C6’s deep intronic variant in *CHM* has been experimentally validated as creating a cryptic splice site [26]. Patients C1, C4 and C8 had not received a genetic diagnosis prior to the long-read sequencing performed in this study. C2 and C3 are sisters, C10 and C11 daughter and mother, R1 and R4 are mother and daughter.

| ID | gene | variant | variant type | impact | grading |
| --- | --- | --- | --- | --- | --- |
| C1 | *CHM* | c.529del; p.Glu177Lysfs*20 | indel | frameshift | 3 |
| C2 | *CHM* | g.85957852del; c.940+3delA | indel | splicing | 4 |
| C3 | *CHM* | g.85957852del; c.940+3delA | indel | splicing | 2 |
| C4 | *CHM* | c.1214_1215insC; p.Gln405Hisfs*13 | indel | frameshift | 2 |
| C5 | *CHM* | c.1780delC; p.Leu594Phefs*55 | indel | frameshift | 2 |
| C6 | *CHM* | g.86045359_86047506del; c.27_49+2125del | SV | deletion | 4 |
| C7 | *CHM* | c.757C>T, p.Arg253Ter | SNV | stop | 3 |
| C8 | *CHM* | g.85838231_85933652del; c.1167-22313_*26400del | SV | truncation | 2 |
| C9 | *CHM* | g.85965588T>C; c.315-1536A>G | SNV | splicing | 3 |
| C10 | *CHM* | c.1584_1587delTGTT; p.Val529Hisfs*7 | indel | frameshift | 2 |
| C11 | *CHM* | c.1584_1587delTGTT; p.Val529Hisfs*7 | indel | frameshift | 1 |
| C12 | *CHM* | g.84295184_84296004del; c.(1770+1_1771-1)_(*1_?)del | SV | truncation | 3 |
| R1 | *RPGR* | c.2362_2366del, p.Glu788Argfs*45 | indel | frameshift | 4 |
| R2 | *RPGR* | c.619+5G>A | SNV | splicing | 4 |
| R3 | *RPGR* | c.2442_2445del, p.Gly817Lysfs*2 | indel | frameshift | 4 |
| R4 | *RPGR* | c.2362_2366del, p.Glu788Argfs*45 | indel | frameshift | 4 |

Long-read sequencing identified disease-causing variants for all patients, including for those where there was no prior genetic diagnostic available. The long reads and chromosome-wide sequencing approach resolved multiple types of events: exonic SNVs, intronic SNVs predicted to affect splicing, and large structural variants removing parts of the gene (Table 1).

All but one of the 16 patients presented at least one sample with a skew below 0.4 (60:40 or greater). For some patients the extent of the skew varied between samples. For example, C3 demonstrated a highly skewed saliva sample (0.10, or 90:10) whereas their buccal swab sample had a less pronounced skew (0.33). However, when two of a patient’s samples were skewed (< 0.4), in all cases they were skewed in the same direction (Fig. 4B).

**Fig. 4.**
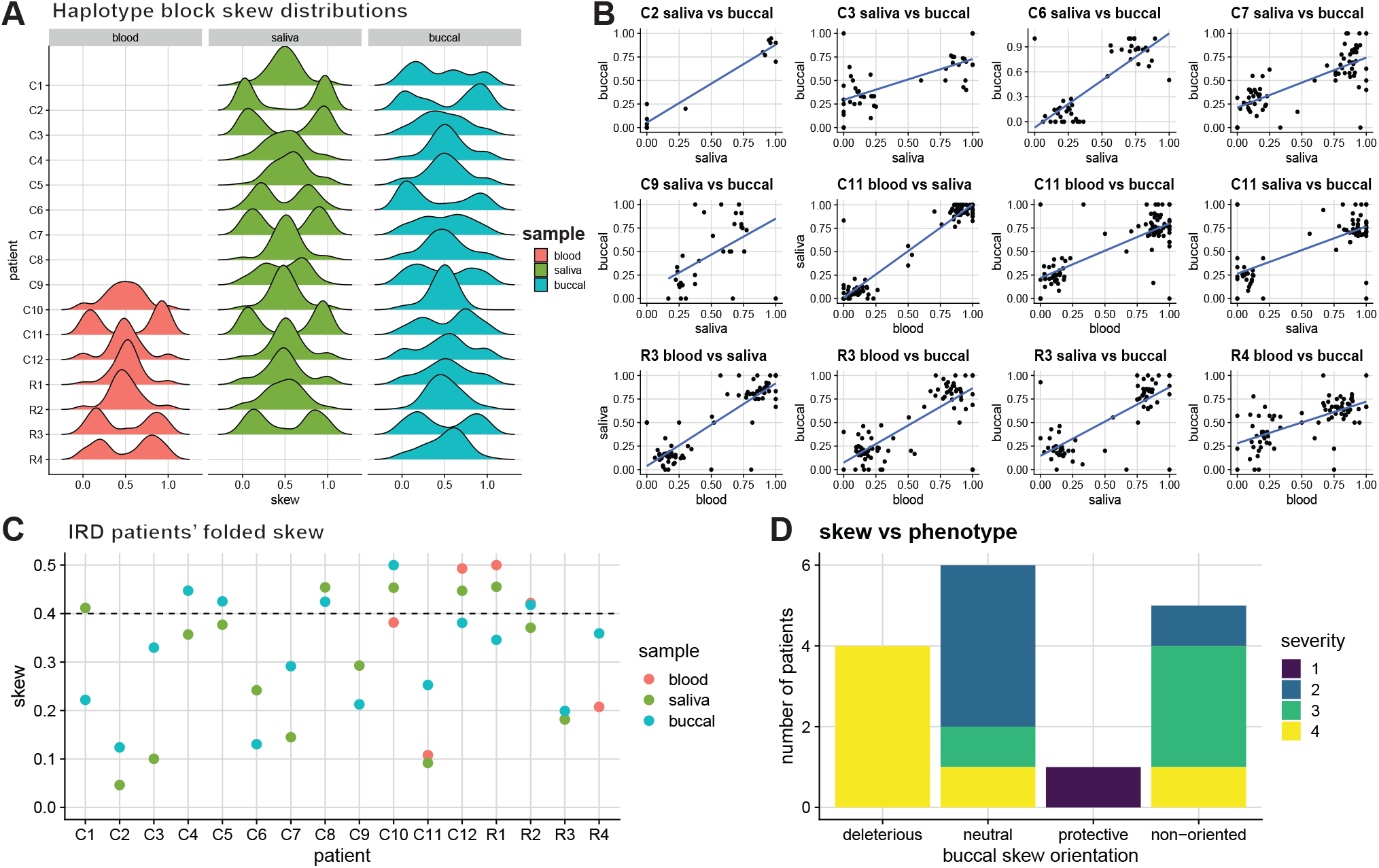
X inactivation skew in a cohort of patients with X-linked inherited retinal disorders. **A)** Distribution of haplotype blocks’ skews in multiple samples from patients carrying mutations in the *CHM* gene (C1-C12) or *RPGR* gene (R1-R4). **B)** Correlations of haplotype blocks’ skew between samples indicates consistency in preferential Xa choice across tissues, for each patient where two or more samples show skew (p < 0.4) **C)** Summary of tissue skews for each patient. **D)** Correlation of buccal skew with phenotypic severity. The skew is classified as either deleterious (variant allele is preferentially Xa), protective (variant allele is preferentially Xi), neutral (no or little skew, p *≥* 0.4). For 6 *CHM* variant carriers with skewed buccal swab samples, the skew could not be oriented (non-oriented category). Retinal severity grading presented in this graph is depicted as grades 1–4 to combine retinal phenotypes for choroideremia and *RPGR*-associated X-linked retinitis pigmentosa [6, 25]: grade 1 is fine and normal phenotypes, grade 2 is coarse and radial pattern phenotypes, grade 3 is geographic and focal pigmentary retinopathy phenotypes, and grade 4 is male pattern phenotypes for choroideremia and *RPGR*-associated X-linked retinitis pigmentosa, respectively.

To correlate skew and phenotype, we used the buccal swab skew as the most likely to reflect the retina’s skew, as it is less sensitive to age-related skewing than blood and saliva [4, 23]. Among the ten severely affected patients (graded as 3 and 4), nine (90%) had a skew of 60:40 or greater, and five (50%) had a skew of 75:25 or greater. Relative to a baseline expectation of 5% of the population with skews 75:25 or greater [4], we concluded that there was an over-representation of skew among affected patients (p-value = 6e-05, one-sided binomial test).

We next oriented the skew relative to the allele that carries the variant: the skew was deemed deleterious when the predominantly active X is the one carrying the variant, and protective when the variant is instead on the predominantly inactive X. For all skewed patients with *RPGR* variants, the skew could be oriented, and all were deleterious (3 out of 3, one patient not skewed). For *CHM* variant carriers however, skew orientation was less successful as CpG islands are more sparse in this genomic region and the length of the haplotype blocks over *CHM* were often limited by stretches of low complexity sequence. Out of 8 skewed patients (4 were not skewed), two could have their skews oriented: one was strongly deleterious (87:13), consistent with the male pattern/grade 4 phenotype, and one was strongly protective (75:25), consistent with the mildest phenotype observed (fine phenotype/grade 1). There-fore out of 5 oriented skews, all correlated with the phenotype (p-value = 0.031, one-sided binomial test). By contrast, patients with a balanced buccal swab X inactivation ratio (neutral skew) were split across phenotypic severity classification: four coarse phenotype (grade 2), one geographic phenotype (grade 3) and one male pattern phenotype (grade 4). These data suggest that a strong deleterious skew measured in peripheral tissues could be predictive, but a neutral skew does not necessarily mean disease will not be severe.

### Investigating the correlation of X inactivation skew correlation and disease severity in patient pairs

We further explored the relationship between choroideremia disease severity and X inactivation skew by focusing on a sister-sister and a mother-daughter pair, each pair sharing the same disease-causing variant as well as 50% of their genetic material (5). In addition to explaining differences in overall disease severity, skewed X inactivation may also be able to explain the geographical patterns of retinal degeneration observed in choroideremia and RPGR-linked retinitis pigmentosa (Fig. 5A and B). In particular, geographic choroideremia and focal pigmentary pattern in retinitis pigmentosa could reflect local patches of cells with elevated deleterious skews, while male patterns may reflect uniformly deleterious skew across the tissue.

**Fig. 5.**
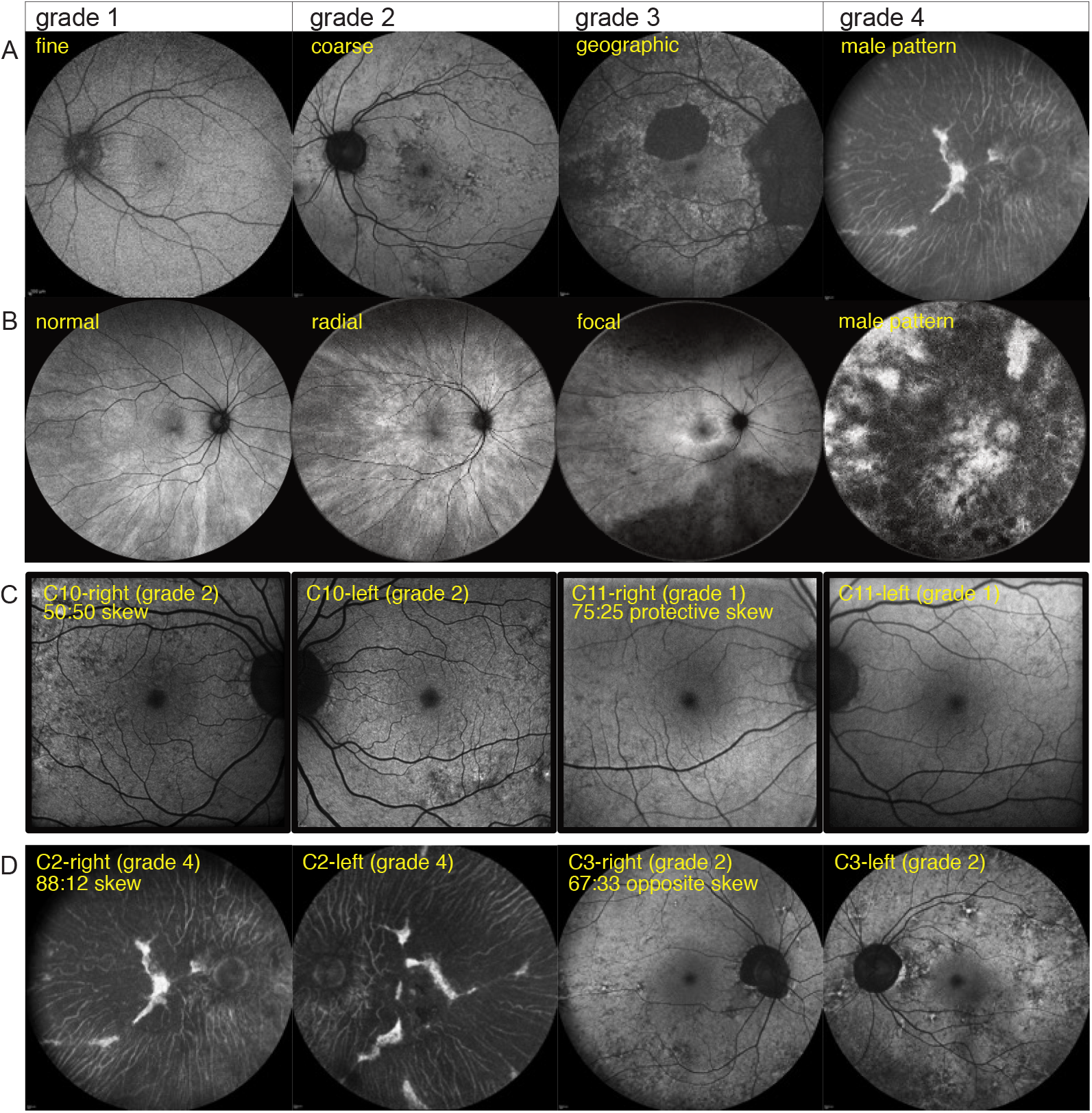
Clinical phenotypes of *CHM* and *RPGR* variant carriers and correlation with X inactivation skew. **A)** Classification of retinal disease in female carriers of choroideremia based on 55-degree retinal fundus autofluorescence images, illustrative of the classification by Edwards et al. [6]: fine, coarse, geographic and male pattern degeneration. **B)** Classification of retinal disease in female carriers of *RPGR*-associated X-linked retinitis pigmentosa based on 55-degree retinal fundus autofluorescence images, illustrating the classification by Nanda et al. [25]: normal, radial pattern, focal pigmentary retinopathy and male pattern degeneration. **C)** Discordant clinical phenotypes of 34-year-old female C10 and her 59-year-old mother C11 carrying the same pathogenic *CHM* variant. C10 presents with coarse (grade 2) phenotype and C11 presents with fine (grade 1) phenotype. C11 had a pronounced buccal swab X inactivation skew (75:25) in favour of expressing the healthy allele, consistent with a milder disease phenotype. 30-degree fundus autofluorescence images of right and left eyes are depicted for both carriers. **D)** Discordant clinical phenotypes of 73-year-old female C2 and her 64-year-old sister C3, carrying the same *CHM* variant. C2 presents with male pattern (grade 4) degeneration and C3 presents with a coarse (grade 2) phenotype. C2 and C3 had skews in opposite directions, but the direction relative to the variant could not be determined. 55-degree fundus autofluorescence images of right and left eyes are depicted for both carriers.

In the first pair, carrying a 4-nucleotide deletion in *CHM* causing a frameshift, the non-skewed (50:50) 34-year-old female C10 presented with coarse (grade 2) phenotype whereas her 59-year-old mother C11 with a strong protective 75:25 skew had a milder fine (grade 1) phenotype (Fig. 5C and D). For a degenerative disease and given the common mutation, the mother’s higher age would be expected to cause a more severe phenotype than her daughter’s; in this case it may have been counteracted by the mother’s skewed X inactivation, favouring the expression of the healthy allele, which provided her with protection against choroideremia. While her daughter is affected, the absence of a strong deleterious skew may be beneficial in avoiding a stronger male pattern phenotype.

In the second pair, a one-nucleotide deletion in the *CHM* coding sequence caused both sisters C2 (73 years old) and C3 (64 years old) to develop choroideremia, but to very different extent (Fig. 5E and F). C2 had a very strong 88:12 skew and a grade 4/male pattern retinal degeneration, suggesting that the skew was deleterious. Unfortunately the *CHM* variant was not contained within a haplotype block with informative CpG islands, so the direction of the skew could not be formally ascertained. However, we could determine that her sister C2 had a skew in the opposite direction (33:67), and therefore protective, which matched with her much milder grade 2/coarse phenotype. These examples thus illustrate the high explanatory power of skewed X inactivation in X-linked diseases.

## Discussion

In this study we demonstrate the use of long-read genomic sequencing to measure skewed X chromosome inactivation. The same data also allows calling of small and large variants: we detected pathogenic intronic and exonic small variants, as well as large structural variants, in a cohort of female patients affected by X-linked retinal disorders. It confirmed previous work outlining the superiority of long reads over short reads to detect variants in the repetitive exon 15 of *RPGR* [26–28] or structural variants in *CHM* [29] among other genes [30]. Where typically multiple assays might be necessary to obtain this information (short-read exome sequencing, short-read genome sequencing, genome CNV microarray, restriction-enzyme/PCR-based tests of skew [31]), long-read sequencing combined with our new skew analysis method has the potential to replace all of those with one workflow, dramatically accelerating the diagnostic journey and reducing its cost. As more and more long-read population genomics projects get underway, factoring in skewed X inactivation (or epigenotype) is likely to improve the genotype-phenotype correlations in females.

Our method takes a chromosome-wide approach, drawing data from hundreds of CpG islands to derive an accurate estimate of the X inactivation skew in the sequenced sample. All samples are informative, contrary to traditional tests based on a single locus that is not sufficiently polymorphic in 20% of cases [11, 13]. By using adaptive sampling to enrich for X-chromosome reads, we obtain a 3–4-fold reduction in sequencing cost per sample while keeping the sample preparation very straightforward (standard ligation-based library preparation protocol). However, our analysis is equally applicable to whole-genome, unbiased sequencing as long as X chromosome coverage is 15 or above. This lower limit on coverage is imposed by the requirement to cluster reads overlapping CpG islands into two groups based on their methylation pattern. With lower coverage, the two groups may not be sufficiently sampled and cluster definition becomes more difficult. It is important to note that skew can vary from tissue to tissue within one individual [4], and certain tissues become more clonal/skewed with age [32, 33]. Therefore there is a risk in extrapolating to a different, inaccessible, disease-relevant tissue. This risk can be mitigated by sampling multiple tissues. In our study we prioritised the buccal mucosa skew to extrapolate to the retina, while the saliva and blood skews could provide additional evidence of a global, organism-wide skew and increase the confidence in the extrapolation.

Orienting the skew relative to the disease-causing variant in the absence of trio data varied in difficulty according to the locus. For a CpG-island rich, repeat-poor locus such as *RPGR*, in all patients (4/4) the haplotype block containing the variant spanned several informative CpG islands and therefore the skew could be oriented with high-confidence without any trio data. By contrast, phasing of the *CHM* locus was often interrupted by complex regions or runs of homozygosity. With the nearest CpG island at the *DACH2* promoter 300 kb away from the 3’ end of the *CHM* coding sequence, for 6 out of 12 patients the haplotype block carrying the variant did not overlap a CpG island. Two of the skews could be oriented using relatedness, but for the remaining patients, trio data would be required to definitively orient the skew. This would not require long-read sequencing but could be done simply by PCR genotyping.

The contribution of X inactivation skew to X-linked inherited retinal disease phenotypes in females remains understudied [8, 9, 34]. In our cohort of 16 affected females, there was an over-representation of skewed X inactivation, and in all severely affected females we found the skew orientation to be deleterious (or in one case predicted to be deleterious with high confidence). By contrast, protective skew in the female relatives of severely affected patients correlated with milder symptoms. Cumulative with previous reports of correlation between skew and *RPGR*-associated retinitis pigmentosa [11], this confirmed the potential for high explanatory power of skewed X inactivation in X-linked inherited retinal disease manifestation in females, and encourages investigating its use in prognostics. As gene therapy solutions for inherited retinal diseases are emerging, including for *RPGR* and *CHM* [35–38], early identification of candidate patients who would benefit from this treatment is crucial, in order to rescue expression before irreversible retinal damage.

Besides assessing the skew for individual variants, quantifying X inactivation skew can also be used to improve phasing of the X chromosome. For strongly skewed samples, haplotype blocks overlapping CpG islands can be linked so haplotype 1 is consistently a single parental allele. Analysis of epiallele bias may also be extended to autosomes, for instance in pharmacogenetic panels, if there is evidence of variable expression correlating with differences in DNA methylation.

In conclusion, reporting X inactivation skew adds an important additional layer of information for all X-linked diseases or risk variants. As such we expect our method to become part of the routine analysis pipeline for long-read population genomic datasets.

## Methods

### Sample collection

This multisite study included two cohorts of female carriers of RPGR-associated retinitis pigmentosa and choroideremia. The Australian site received ethics approval from The Royal Victorian Eye and Ear Hospital Human Research Ethics Committee (Ethics ID: 19-1443H). The United Kingdom (UK) site received ethics approval from the Anglia Ruskin University Human Research Ethics Committee (Ethics ID: ETH2223-5360). The wet lab component of this study received ethics approval from the Walter Eliza Hall Institute (Ethics ID: 20/16B). Written informed consent was obtained from each participant prior to commencement of the study, which was undertaken at the respective sites.

We used Oragene OG-600 and OCR-100 kits (DNA Genotek) for saliva and buccal mucosa collection, respectively. Patients provided saliva samples by spitting into a tube until the saliva (not foam) passed the fill-line. Buccal swab involved using the cotton tip in the collection kit to rub inside the cheek (no scraping involved). The genetic variants for the participants are outlined in Table 1.

### Clinical visits

The clinical component of this study was part of a larger project investigating retinal phenotypes of female carriers of X-linked inherited retinal disorders, which involved participants attending a research visit for a comprehensive ocular examination. Of interest to the current study, participants had fundus autofluorescence imaging performed to classify their retinal severity according to current grading scales for RPGR-associated retinitis pigmentosa [25] and choroideremia [6], and a selection of these images have been utilised in this publication.

### DNA extraction

Blood, saliva and buccal mucosa samples underwent DNA extraction at the Australian Genome Research Facility (AGRF) using a QIAgen DNeasy blood & tissue kit. Retinal DNA was extracted from 3 mm diameter retinal discs with a QIAamp DNA Micro Kit.

### Nanopore library preparation

Extracted genomic DNA was quantified with a Qubit and assessed for purity on a Nanodrop and assessed for fragmentation on a TapeStation. Samples that showed excessive fragmentation were size selected in one of two ways: when more than 3 µg of DNA were available, the samples were subjected to semi-selective DNA precipitation (v2, https://community.nanoporetech.com/extraction_methods/size-selection2) based on Jones et al. [39]); if fewer than 3 µg were available, we used a 1X High-Molecular Weight bead clean-up [39]. 1 µg of DNA for each sample was then sheared to a 10–20 kb target range using Covaris G-tubes (two 30-second spins at 3,000 g) and taken forward for library preparation following the instructions for genomic DNA sequencing with barcoding kit 14 (SQK-NBD114.24, Oxford Nanopore Technologies). Three-to-four barcoded samples were pooled together for each library to be sequenced on one flow cell.

### Nanopore sequencing

160 ng of library were loaded on R10.4.1 PromethION flow cells (FLO-PRO114M), and run to exhaustion on a PromethION P24 running MinKNOW v23.04.6, using adaptive enrichment of the X chromosome from the T2T-CHM13v2 genome reference.

### Adaptive sampling calibration

We start by deriving an estimate of enrichment efficiency as a factor of read lengths, to inform library preparation strategy. When enriching for a whole chromosome or in cases where the target length is much larger than read lengths (*L*_target_ *≫ L*_read_), on-target output is maximised by having the longest read lengths possible. As reads are sampled at random, without adaptive sampling the proportion of the time *p* that a pore spends acquiring data is simply proportional to the target space (*p ∝ L*_target_*/L*_genome_) and is independent of read lengths if the wait time between reads *t*_wait_ is much smaller than the time it takes to sequence a read *t*_seq_ (*t*_wait_ *≪ t*_seq_). However with adaptive sampling, on average the pore has to sample *n*_off_ = *L*_genome_*/L*_target_ off-target reads before finding a read that is on-target. So the proportion of time spent acquiring on-target data is reduced to

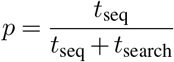

where *t*_search_ is the time spent trying and discarding off-target reads, composed of *n*_off_ repeats of *t*_reject_ + *t*_wait_, with *t*_reject_ the time it takes to sample and reject an off-target read. By denoting the sequencing speed 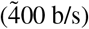 as *s*, we have 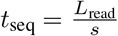, and thus

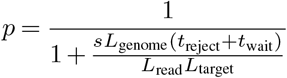

In the case of a human genome (3.1 Gb) and targeting the entire X chromosome (154 Mb), taking *t*_reject_ = 2.3 s based on rejected reads of length 900 b and assuming *t*_wait_ = 0.5 s, with 1 kb reads the proportion of time spent acquiring on-target data is only *p* = 4.2% while with 10 kb reads it increases to 31%.

### Skew analysis pipeline

Reads from POD5 files were basecalled and mapped to the T2T-CHM13v2 human genome reference using dorado v0.4.2 (https://github.com/nanoporetech/dorado), which uses minimap2 [40]. Modified basecalling was performed concurrently for 5mCG using the dorado argument `--mod_bases 5mCG`. Mosdepth v0.3.3 [41] and NanoComp v1.21.0 [42] were used for coverage calculations and quality control. Modbam files from each patient’s samples were merged to call SNVs with DeepVariant v1.5.0-gpu [18] on the X chromosome using the ONT_R104 model, and filtered for high-quality ‘PASS’ calls using bcftools [43]. The variant call file (VCF) was then phased using WhatsHap v1.7 [19] using the merged modbam file. Each sample modbam (saliva, buccal, and blood and retina if available) was then processed individually by NanoMethViz v2.7.8 [20] to cluster reads by their CpG island methylation pattern, using the HDBSCAN algorithm [44] and setting the minimum number of reads to form a cluster as 5. The UCSC CHM13v2 (or mm10 for mouse) CpG island annotation was used. CpG islands for which there were exactly two clusters were used to assign Xa and Xi labels to the reads of each cluster (Xa for the low methylation cluster, Xi for the high methylation cluster, reversing these labels in the case of the *XIST/Xist* promoter CpG island). Within each haplotype block, the number of haplotype 1/Xa, haplotype 2/Xi, haplotype 1/Xi and haplotype 2/Xa reads were counted to estimate the block-wise skew, and to estimate the sample skew as the value that maximises the joint probability of observing those counts over all blocks.

Given this sample skew, log-odds of haplotype 1 being the predominantly active vs inactive X for a given haplotype block is given by logodds = (2 *∗* successes *−* trials) *∗ log*10(*p*) + (trials *−* 2 *∗* successes) *∗ log*10(1 *− p*).

We used *R* packages *plyranges* v1.20.0 [45] for genomic interval manipulation and cowplot v1.1.1 for visualisation.

## Data availability

The mouse neural stem cells ultra-long nanopore gDNA sequencing from Su et al. [17] is available under accession E-MTAB-13706. The patients’ data cannot be made public under our ethics approvals.

## Acknowledgements

We thank Sarah MacRaild, Stephen Wilcox and Marek Cmero for their assistance running the PromethION, and Jafar Jabbari for sharing his custom bead clean-up protocol.

## Supplementary information

**Extended Data Fig. 1.**
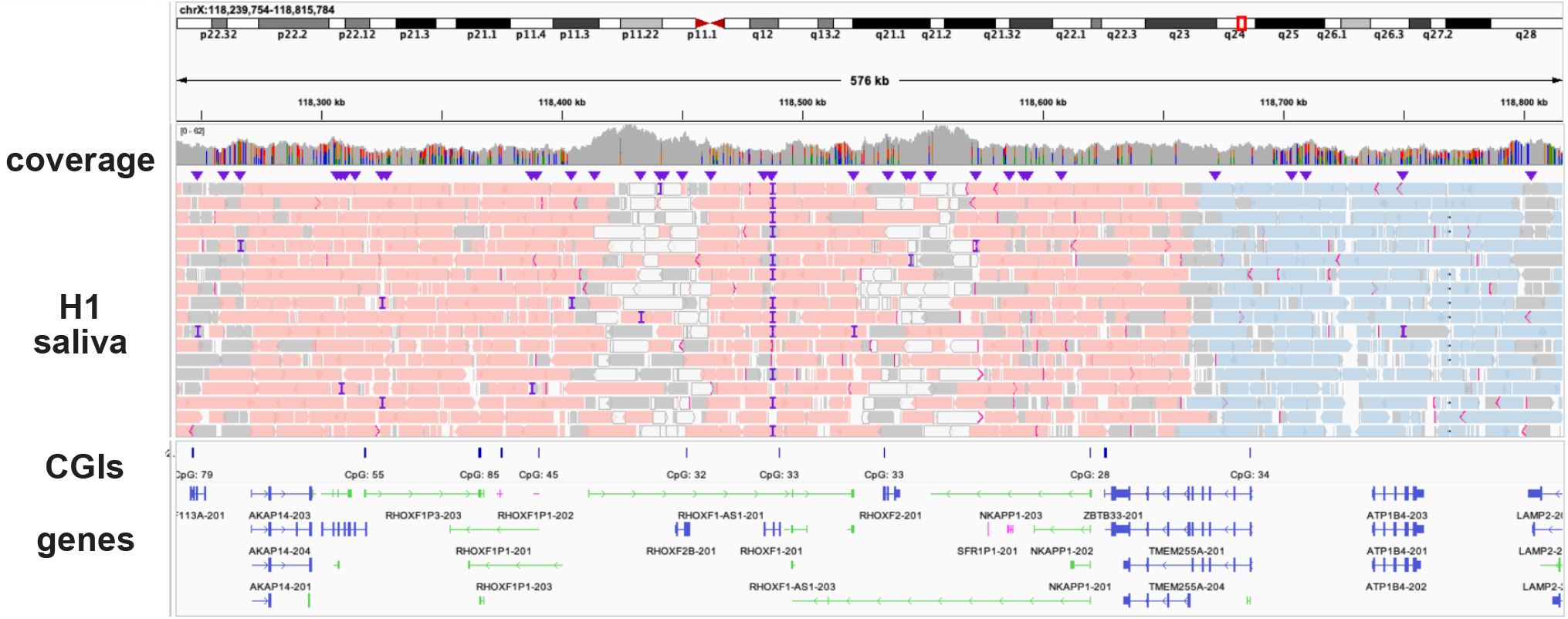
Skew analysis identifies a phasing error. IGV screenshot of H1 saliva sample reads mapped to the *RHOXF2*/*RHOXF2B* region. Reads are coloured by haplotype block. The tandem repeat leads to ambiguous mapping (white reads) and a phase-switching error within the red haplotype block.

